# Strong, but not weak, noise correlations are beneficial for population coding

**DOI:** 10.1101/2024.06.26.600826

**Authors:** Gabriel Mahuas, Thomas Buffet, Olivier Marre, Ulisse Ferrari, Thierry Mora

## Abstract

Neural correlations play a critical role in sensory information coding. They are of two kinds: signal correlations, when neurons have overlapping sensitivities, and noise correlations from network effects and shared noise. It is commonly thought that stimulus and noise correlations should have opposite signs to improve coding. However, experiments from early sensory systems and cortex typically show the opposite effect, with many pairs of neurons showing both types of correlations to be positive and large. Here, we develop a theory of information coding by correlated neurons which resolves this paradox. We show that noise correlations are always beneficial if they are strong enough. Extensive tests on retinal recordings under different visual stimuli confirm our predictions. Finally, using neuronal recordings and modeling, we show that for high dimensional stimuli noise correlation benefits the encoding of fine-grained details of visual stimuli, at the expense of large-scale features, which are already well encoded.

## I. INTRODUCTION

Neurons from sensory systems encode information about incoming stimuli in their collective spiking activity. This activity is noisy: repetitions of the very same stimulus can drive different responses [1–6]. It has been shown that the noise is shared among neurons and synchronizes them, an effect called *noise correlations*, as opposed to *signal correlations* induced by the stimulus [6– 10]. Noise correlations have been observed since the first synchronous recordings of multiple neurons [11, 12] and at all levels of sensory processing, from the retina [2, 13– 21] to the visual cortex [1, 4, 5, 22–25] and other brain areas [6, 7, 9, 26–29]

Strong noise correlations have been measured mostly between nearby neurons with similar stimulus sensitivity [1, 5, 12, 23, 27, 28, 30–32]. This behaviour is particularly evident in the retina between nearby ganglion cells of the same type [2, 14, 16, 17, 21]. This observation is however surprising, since previously it was thought that these correlations are detrimental to information coding: a theoretical argument [1, 33–36] suggests that noise correlations are detrimental to information transmission if they have the same sign as signal correlations [7, 9, 10]. This rule is sometimes called the sign rule [37], and is related to the notion of information-limiting correlations [38]. Since nearby neurons with similar tuning are positively correlated by the signal, the theory would predict that their positive noise correlations should be detrimental, making the code less efficient. However, a large body of literature has reported the beneficial effects of noise correlations on coding accuracy [15, 18, 20, 39–44]. Because of these contradictions, the effect of shared variability on information transmission is still unclear, and remains a largely debated topic in neuroscience [7–9].

Here we aim to resolve these tensions by developing a general framework that builds on previous theoretical work [45] and is grounded on the analysis of multi-electrode array recordings of rat and mouse retinas. While previous studies have considered the impact of noise correlations either for particular stimuli [1, 15, 18, 29], or for particular models [36, 40, 41], our approach is general and covers both low and high dimensional stimuli. We show that the sign rule can be broken in a specific regime that we observed in retinal responses: when noise correlations are strong enough compared to signal correlations, they have a beneficial effect on information transmission. Our results unravel the complex interplay between signal and noise correlations, and predict when and how noise correlations are beneficial or detrimental. In the case of high dimensional stimuli, like images or videos, our theory predicts different effects of noise correlations depending on stimulus features. In particular, it explains how large noise correlations between neurons with similar stimulus sensitivity help encode fine details of the stimulus.

We study theoretically the different regimes for pairs of spiking neurons, and illustrate them in the correlated activity of rat retinal ganglion cells. We then extend our analysis to large populations of sensory neurons, and propose a spectral analysis suggesting that local noise correlations enhance information by favoring the accurate encoding of fine-grained details. We validate this last prediction combining data from the mouse retina with accurate convolutional neural network (CNN) models.

## II. RESULTS

### Strong pairwise noise correlations enhance information transmission

We start with a simple model of a pair of spiking neurons encoding an angle *θ*, for instance the direction of motion of a visual stimulus, in their responses *r*_1_ and *r*_2_. These responses are correlated through two sources: signal correlations *ρ*_s_ due to an overlap of the tuning curves (Fig. 1A); and noise correlations *ρ*_n_ due to shared noise (see Methods for mathematical definitions). We asked how this shared noise affects the encoded information, for a fixed level of noise in neurons.

**FIG. 1:**
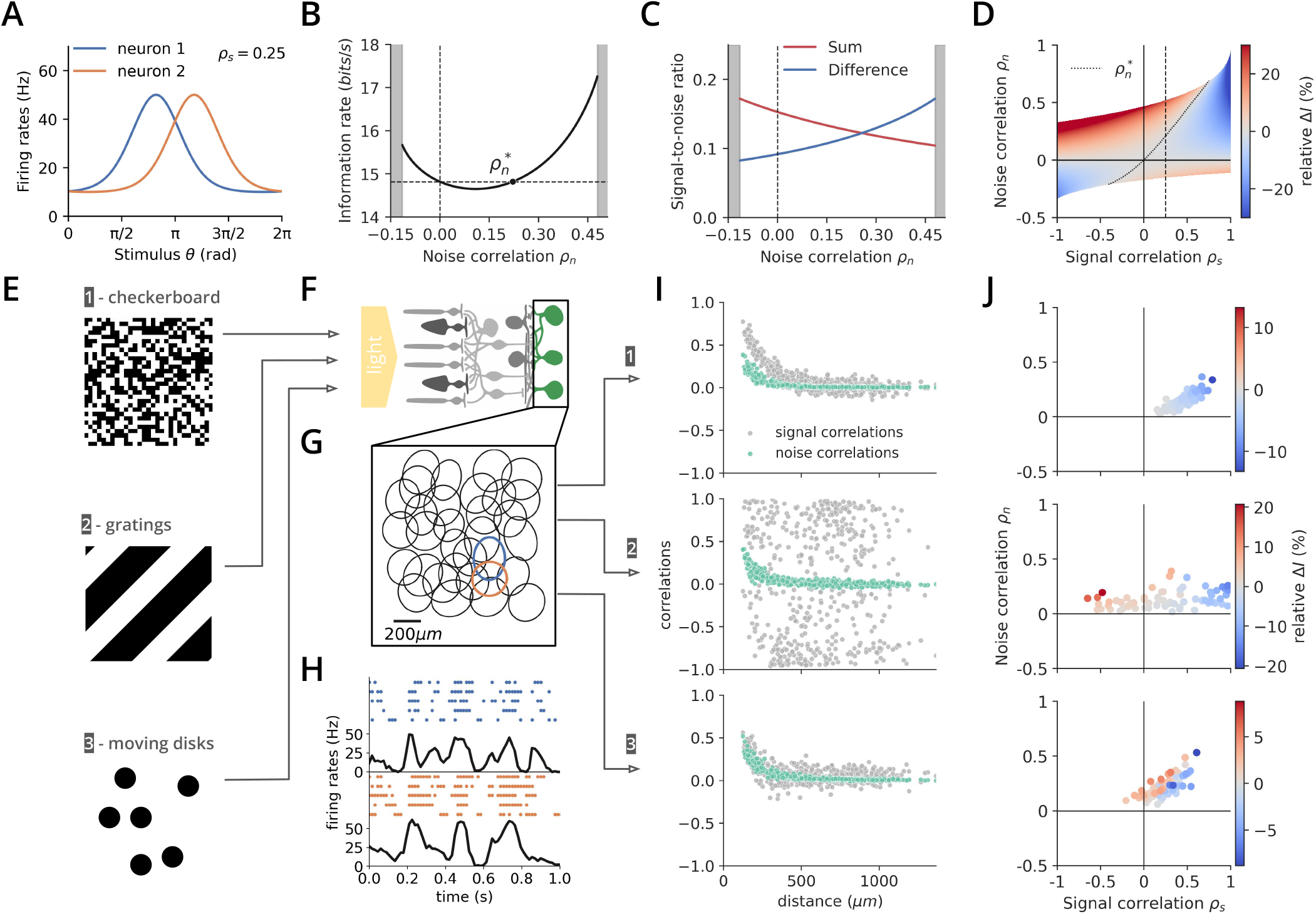
The effects of noise correlations on information coding depends on the stimulus. **A**. Example pair of Von Mises tuning curves with moderate signal correlation level (*ρ*_*s*_ = 0.25). **B**. Mutual information between stimulus and response for the example pair of A, vs the strength of noise correlations. Grey areas correspond to forbidden correlations zones. **C**. The non-monoticity of B may be explained by examining how well the stimulus is represented by the sum and difference of the two neurons’ activities, as measured by their signal-to-noise ratios. Noise correlations enhance noise in the sum, but reduce it in the difference. **D**. Heatmap representing the noise synergy, defined as the relative gain of mutual information induced by noise correlations compared to the uncorrelated case. The dotted vertical line corresponds to the example pair of A and B. **E**. Three stimuli with different spatiotemporal statistics were presented to a rat retina. **F**. Retinal ganglion cells (RGCs) were recorded using a multi-electrode array (MEA). **G**. We isolated a nearly complete population of OFF-*α* cells, with receptive fields (RFs) that tile the visual field following approximately a triangular lattice. **H**. Example raster plots and firing rates for two cells with neighboring RFs. **I**. Signal and noise correlations for each pairs of neurons in the population, versus their distance. Each plot corresponds to 1 of the 3 stimuli of E. **J**. Noise synergy induced by noise correlations for all pairs of nearby neurons (≥ 300*µm*), for each stimulus of E.

To quantify the joint coding capacity of the 2 neurons, we computed the mutual information *I*(*θ*; *r*_1_, *r*_2_) between their activities and the stimulus *θ*. For fixed tuning curves, we find that the mutual information depends non monotonously on the noise correlation *ρ*_n_ (Fig. 1B). For small abolute values of *ρ*_n_, the sign rule is satisfied, meaning that negative noise correlations are beneficial, and weak positive ones are detrimental [7, 35, 37, 39]. However, the mutual information increases again and noise correlations become beneficial if they are larger than a certain threshold 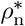, violating the sign rule. This non monotonous dependency may be intuitively explained as the interplay between two opposite effects (Fig. 1C). Negative noise correlations are beneficial because they reduce noise in the total activity of the neurons. By contrast, positive noise correlations reduce noise in their differential activity, but this effect only dominates when they are strong enough.

We call “noise synergy” the gain in information afforded by noise correlations, Δ*I* = *I*(*ρ*_n_) ™ *I*(*ρ*_n_ = 0). Fig. 1D shows how noise synergy depends on both the noise and signal correlation, where the latter is varied in the model by changing the overlap between the tuning curves. Very generally, and beyond the cases predicted by the sign-rule, noise correlations are beneficial also when they are stronger than the signal correlations. We can gain insight into this behaviour by computing an approximation of the mutual information that is valid for small correlations, following [45] (see Methods). The noise synergy can be expressed as:

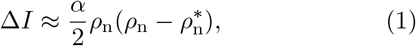

where *α ≤* 1 is prefactor that grows with the signal-to-noise ratio (SNR) of the neurons. Eq. 1 captures the behaviour of Fig. 1B, in particular the observation that noise correlations are beneficial if *ρ*_n_*ρ*_s_ < 0, as the sign rule predicts, or if they are strong enough, 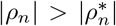. We can show (see Methods) that the threshold 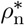 scales with the signal correlation strength *ρ*_s_:

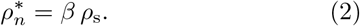

This result holds in the case of Gaussian neurons (see Methods) and the prefactor *β ≤* 1 gets smaller and even approaches 0 as the SNR increases. It is also smaller when these SNR are dissimilar between cells, consistent with previous reports [41]. When the SNRs are weak and similar, we have *β* ≈1. This analysis indicates that noise correlations are beneficial when they are of the same strength as signal correlations, but also that this benefit is enhanced when neurons are reliable.

Our definition of the noise synergy relies on comparing the noise-correlated and uncorrelated cases at fixed noise level or SNR. However, increasing noise correlations at constant SNR decreases the effective variability of the response, as measured by the noise entropy of the joint response of the pair (see Methods). This means that high noise correlations imply a more precise response, which could explain the gain in information. To study this possible confounding factor, we also computed Δ*I* at equal noise entropy, instead of equal SNR, and found that strong noise correlations are still beneficial, with modified 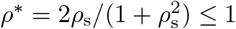 (see Methods).

### Benefit of noise correlations in pairs of retinal ganglion cells

The theory predicts that noise correlations may be beneficial when they are of the same sign and magnitude as the signal correlations. To see whether real neurons fall in that physiological regime, we recorded *ex vivo* the joint spiking activity of rat retinal ganglion cells (RGCs, see Methods). We subjected the same retinal preparation to 3 stimuli with distinct spatio-temporal patterns: a random flickering checkerboard, drifting gratings, and randomly moving disks (Fig. 1E). The activity of RGCs was recorded using a multi-electrode array (Fig. 1F), and data was processed to assign spikes to each neuron [46]. We identified cells belonging to a nearly complete OFF-*α* population forming a regular mosaic pattern of their receptive fields (Fig. 1G).

Each of the 3 stimulus movies was repeated multiple times (Fig. 1H), which allowed us to compute the noise and signal correlation functions *ρ*_n_ and *ρ*_s_ (Fig. 1I), see Methods. All three stimuli produced similar structures of noise correlations across the network, with positive correlations between cells with nearby receptive fields. This is consistent with the fact that noise correlations are a property of the network, independent of the stimulus [21, 47], and likely come here from gap junctions coupling neighbouring RGCs [16, 48]. In contrast, signal correlations strongly depend on the statistical structure of the presented stimulus, and may be positive or negative, with varying strengths.

To test the predictions of our theory, we computed the mutual information between stimulus and response for all pairs of cells whose receptive fields were closer than 300 *µ*m (Fig. 1J). The case of the drifting gratings with fixed orientation offers an illustration of the sign rule. That stimulus induces strong negative signal correlations between many cells, depending on their relative positions relative to the gratings direction. Since noise correlations are positive, they are of opposite sign and therefore beneficial. In the case of the checkerboard stimulus, noise correlations were found to be generally detrimental. This again agrees with the sign rule since they have the same sign as signal correlations, but are too weak to surpass the critical correlation value 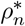. Finally, the case of the moving disks provides an example of the third regime, which violates the sign rule: noise correlations are of the same sign as the signal correlations, but also of comparable magnitude. As a result, many pairs fall above the threshold 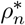, making noise correlations beneficial.

Overall, the 3 stimuli illustrate the 3 possible regimes predicted by the theory when noise correlations are positive: a beneficial effect when signal correlations are negative, a detrimental effect if signal correlations are positive and noise correlation weaker, and a beneficial effect when noise and signal correlations are both positive and of the same magnitude.

### Large sensory populations in high dimension

We then asked how these results extend from pairs to large populations, by considering a large number of neurons tiling sensory space (Fig. 2A). To go beyond neurons tuned to a single stimulus dimension, and account for the ability of neurons to respond to different stimuli in a variety of natural contexts, we assume that each neuron responds to high-dimensional stimulus, like a whole image, a temporal sequence, or a movie. As different stimuli are shown, the spike rate of each neuron will vary. For computational ease, we take these fluctuations to be Gaussian of variance *V*_s_.

**FIG. 2:**
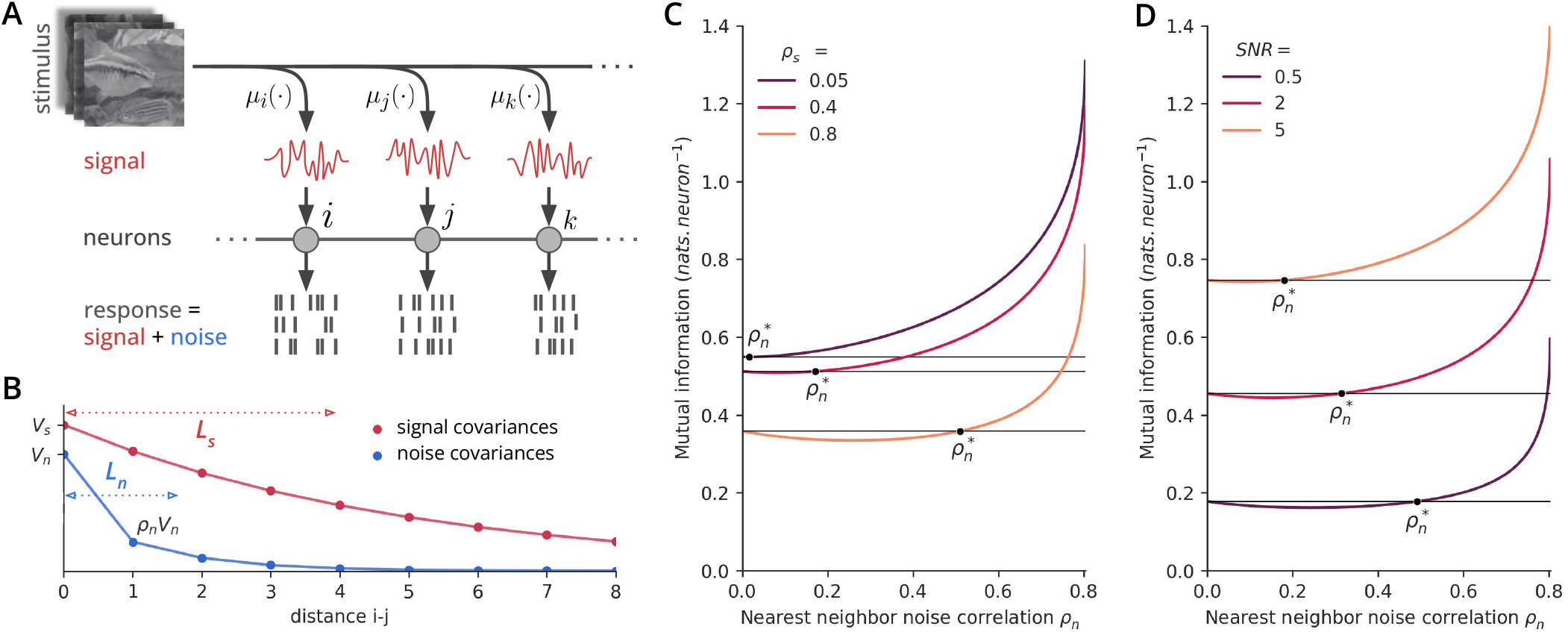
Population analysis. **A**. Neurons are assumed to be spatially arranged along sensory space. They combine features of the stimulus through a response function *µ*_*i*_. Noise is added to the neural responses. **B**. Signal and noise covariances versus distance between neurons. Signal and noise covariances decay exponentially with distance with spatial scales *L*_s_ and *L*_n_. **C.-D**. Mutual information as a function of the noise correlation between neighbors for: (**C**) varying levels of signal correlations, with fixed *V*_s_ = 2, *V*_n_ = 1, and *L*_n_ = 2; and (**D**) varying levels of signal-to-noise ratio (SNR= *V*_s_*/V*_n_), with *L*_s_ = *L*_n_ = 2 (*ρ*_s_ ≈ 0.6).

To account for the empirical observation that nearby neurons tend to have close receptive fields, we correlate the responses of any two neurons with a strength that decreases as a function of their distance in sensory space, with characteristic decay length *L*_s_ (Fig. 2B). The value of the correlation between nearest neighbours quantifies the signal correlation, *ρ*_s_. For simplicity the response noise is also assumed to be Gaussian of variance *V*_n_. To model positive noise correlations between nearby neurons observed in both the retina [2, 14, 16, 17, 21] and cortex [1, 5, 12, 23, 27, 28, 30–32], we assume that they also decay with distance, but with a different length *L*_n_ (Fig. 2B). The noise correlation between nearest neighbors, defined as *ρ*_n_, quantifies their strength.

In this setting, both signal and noise correlations are positive, and the sign rule alone would predict a detrimental effect of noise correlations. The mutual information can be computed analytically in terms of simple linear algebra operations over the neurons’ covariance matrices (see Methods) [49]. Using these exact formulas, we examined how the mutual information changes as a function of the noise correlation *ρ*_n_ for different values of the signal correlation *ρ*_s_ (Fig. 2C) and of the SNR *V*_s_*/V*_n_ (Fig. 2D).

The results qualitatively agree with the case of pairs of neurons considered previously. Weak noise correlations impede information transmission, in accordance with the sign rule. However, they become beneficial as they increase past a critical threshold 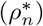, and this threshold grows with the signal correlation strength. It also decreases and even vanishes as the SNR is increased (Fig. 2D and Methods for a discussion of the large SNR limit). This means that more reliable neurons imply an enhanced benefit of noise correlations. We further proved that, even at low SNR, there always exists a range of noise correlation strengths where noise correlations are beneficial (see Methods). The general dependency of 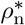 on the correlation ranges *L*_s_ and *L*_n_ is shown in Fig. S1. Based on the analysis of pairs of neurons, we expect inhomogeneities in the SNR *V*_s_*/V*_n_ of neurons to enhance the benefit of noise correlations. To study this effect, we let the power of the signal *V*_s_ vary between cells, while the noise level *V*_n_ is kept constant. Assuming that each cell is assigned a random value of *V*_s_, we can compute the correction to the critical noise correlation 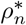. We find that 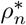 decreases at leading order with the magnitude of the inhomogeneity (see Methods). This result confirms that, in large populations of neurons as well, variability among neurons makes it more likely for noise correlations to have a beneficial effect.

### Spectral decomposition

Mutual information is a single number that provides a global quantification of coding efficiency, but says nothing about what is being transmitted. Likewise, a positive noise synergy indicates that noise correlations are beneficial overall, but it doesn’t tells us what feature of the stimulus are better encoded, nor which specific interactions between signal and noise allow for that benefit. We wondered what features of the signal were enhanced by strong positive noise correlations in our population encoding model.

Thanks to the translation-invariant structure of the model, the mutual information and noise synergies may be decomposed spectrally as a sum over spatial frequencies *k* (expressed in units of inverse distance between nearest neighbors):

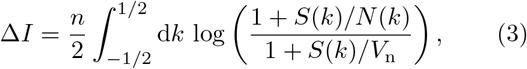

where *S*(*k*) is the power spectrum of the stimulus, and *N* (*k*) that of the noise (see Methods), and *n* → ∞ is the total number of neurons. In this decomposition, low frequencies correspond to long-range collective modes, while high frequencies correspond to fine-grain features.

Natural stimuli involve spatially extended features impacting many neurons. This causes neural responses to exhibit strong long-range signal correlations between neurons, corresponding in our model to large *L*_s_ (Fig. 2B). Most information is then carried by low frequency modes of the response (Fig. 3A).

**FIG. 3:**
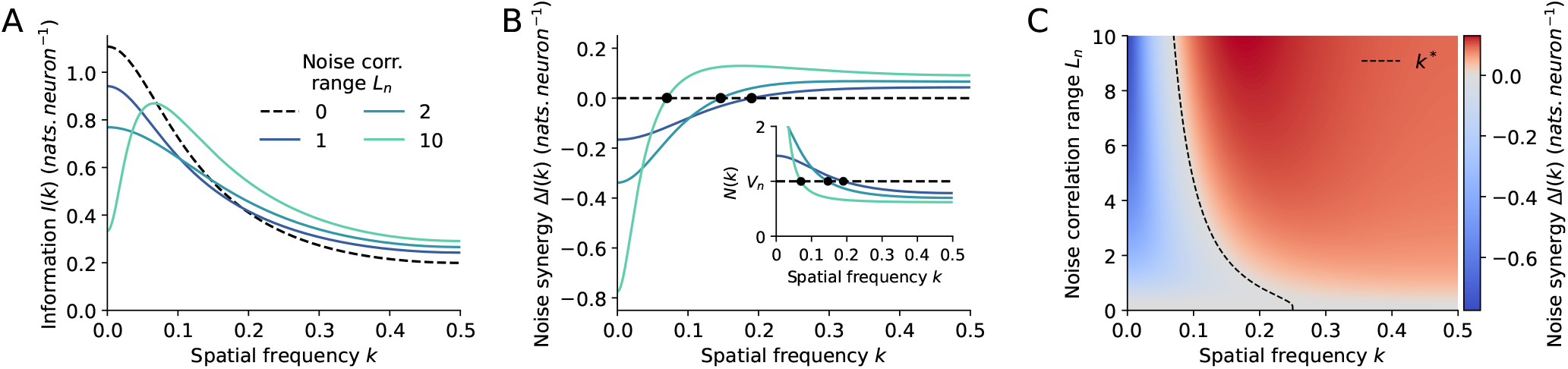
Spectral analysis of stimulus information encoding. **A**. Spatial spectral decomposition *I*(*k*) of the mutual information between stimulus and response for a system with SNR= 2, *L*_s_ = 2 and *ρ*_n_ = 0.4, for various ranges of the noise correlations (*L*_n_ = 0 corresponds to the absence of noise correlations). **B**. Spectral decomposition of the noise synergy Δ*I*(*k*) = log[(1 + *S*(*k*)*/N* (*k*))*/*(1 + *S*(*k*)*/V*_n_)]. The inset shows the power spectrum of the noise. **C**. Heatmap showing the noise synergy spectral decomposition as a function of the noise correlation range *L*_n_. The critical spatial frequency *k*^*^ above which noise correlations are beneficial is shown as a black dotted line.

Noise correlations concentrate noise power at low frequencies and decrease noise power at high frequencies for a fixed noise level *V*_n_ (inset of Fig. 3B). As a result, noise correlations enhance information in the high frequency modes of the signal (*k* ≥ *k*^*^), at the expense of the low frequencies features (Fig. 3B), which are already well represented. Fig. 3C shows the spectral decomposition of the noise synergy as a function of the noise correlation range *L*_n_. The critical frequency *k*^*^ = (1*/*2*π*) arccos 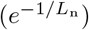 above which noise correlations are beneficial only depends on *L*_n_ (Fig. 3C). However, the relative information gains in each frequency domain depends on the strengths of the signal and noise correlations.

In summary, noise correlations enhance fine details of the stimulus to the detriment of its broad features, which are already sufficiently well encoded. This redistribution of the noise across the spectrum drives the gain in information. This effect is generic to any choice of the correlation lengths, and we expect it to hold for other forms of the power spectra and receptive field geometries.

### Noise correlations in the retina favor the encoding of fine stimulus details

To test our predictions, we studied experimentally the impact of noise correlations on the encoding of features at different spatial scales in the retina. We recorded *ex vivo* the spiking activity of 7 OFF-*α* retinal ganglion cells from a mouse retina using the same experimental technique as described before. We presented the retina with a multiscale checkerboard stimulus composed of frames made of random black and white checkers, flashed at 4 Hz. Each frame was made of a checkerboard with a given spatial resolution (checks of sizes 12, 24, 36, 72 and 108 *µ*m). From the recorded activity, we infered a deep generalized linear model [47] and used the inferred model to build a large synthetic population of 49 cells organized on a triangular lattice (Fig. 4A). We then generated a large dataset of repeated responses to regular black and white checker flashes. Each checker was composed of checks of a given size (sizes ranging from 140 to 420 *µ*m, with 28 *µ*m increments) and for each check size, 50 spatially offset versions of the checker were showed. We trained a linear decoder of each pixel value (black or white) on this synthetic dataset, and a second decoder on the synthetic data in which the activity of each cell was shuffled across repetitions to destroy noise correlations (see Methods).

**FIG. 4:**
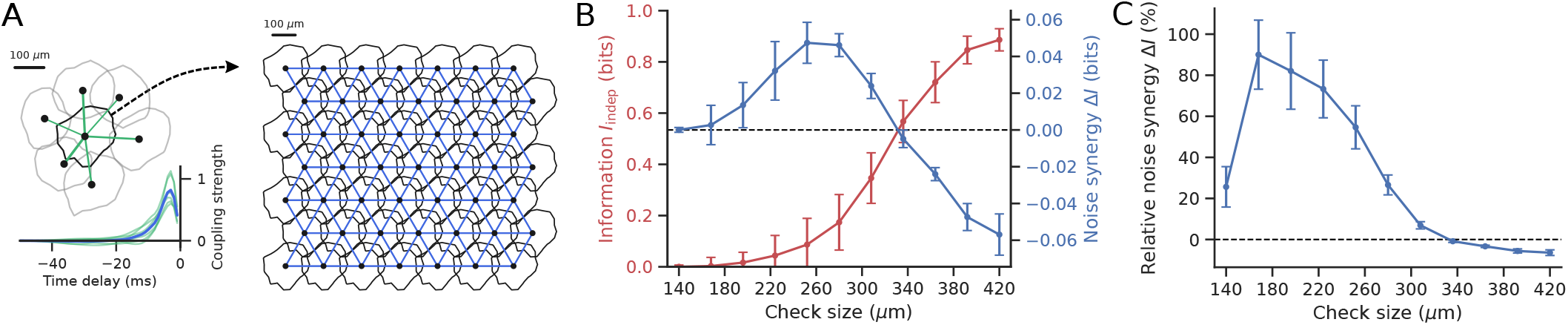
Noise correlations benefit small scale features at the detriment of large scale ones. **A**. We built a large population of 49 RGCs based on 7 neurons recorded from the mouse retina. A deep-GLM [47] was fit to the experimental population and its central neuron model was tilled on a triangular lattice to create a large RGC network. Couplings between the central experimental cell and its neighbors were symmetrized (green links in the population plot; green lines in the inset plot) and averaged to obtain the coupling filters between nearest neighbors in the synthetic population (blue links in the synthetic mosaic; blue line in the inset plot). **B**. Information in the absence of noise correlations *I*_indep_ and noise synergy Δ*I* per pixel for stimulus features of increasing scales. These quantities were computed via a decoding approach applied to a binary flashed checkerboard stimuli with various check sizes. Error bars are the standard error obtained by repeating the analysis on bootstrapped data. In the absence of noise correlations, little information is transmitted about small stimulus features. By contrast, large scale features are well encoded and information per pixel saturates towards 1 bit as check size grows. The noise synergy is positive for small and intermediate check sizes while negative for larger checks, in line with the theoretical results highlighted in Fig. 3. **C**. Noise correlations nearly double the amount of information encoded about stimulus features of small and intermediate sizes, while only decreasing information for the largest checks by less than 10%.

The two decoders were then applied to the testing datasets, synthetically generated in the same way as the training sets, to decode each pixel from the response. For a fair comparison, the second decoder was applied to data in which noise correlations were removed by shuffling, as in the training. The mutual information carried by the decoders was then estimated separately for each checker size. To limit border effects, the mutual information was estimated for each pixel within a small hexagon centered on the central cell of the synthetic population, of size (distance between opposite sides of the hexagon) equal to the distance between cells.

We found that the gain in mutual information afforded by noise correlations is large and positive for small and intermediate check sizes, while moderately negative for large checks (Fig. 4B and C). These results suggest that noise correlation benefit the encoding of small-scale features of the stimulus, at the expense of the large-scale ones, which are easier to encode. Noise correlations can therefore trade the encoding power of large-scale features to improve sensitivity to the small-scale ones.

## III. DISCUSSION

Many experimental works have shown that neurons with the strongest positive noise correlations are similarly tuned to the stimulus [1, 5, 12, 23, 27, 28, 30–32]. Here the sign rule [7, 9, 37] would predict a detrimental effect of shared variability, at odds with the efficient coding hypothesis [50], which is supported by a large body of work showing that noise correlations are indeed beneficial [15, 18, 20, 39, 41, 43]. Our work resolves this inconsistency by showing that beyond a critical value 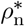, noise correlations become beneficial to information encoding regardless of their sign. We experimentally demonstrated this effect in recordings of retinal neurons subject to stimuli with different statistics, and showed that it generalizes to large populations of sensory neurons.

Pairwise correlations build up to strong network effects for large populations [51]. This large scale synchronization should be detrimental for coding because it impedes denoising by pooling the signal of multiple neurons [1, 36]: the information gain saturates compared to a population of independent neurons. In contrast, other studies focusing on the stimulus response of large sensory populations have observed a positive gain [15, 18, 20, 41, 42]. Our study proposes a solution to this dispute: when the neural population encodes a low dimensional stimulus, as the angle of a drifting gratings, similarly tuned nearby neurons become strongly signal-correlated, and their noise correlations are detrimental [36]. In the case of high dimensional stimuli, like naturalistic images or videos, signal correlations between them are positive but weak, so that noise correlations become larger than the threshold 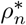 and therefore beneficial. We analyzed the impact of shared variability depending of the stimulus spatial frequency: large scale (low dimensional) modes give rise to strong signal correlations, making positive noise correlations detrimental, while small scale (high dimensional) modes benefit from positive signal correlations since their signal correlations are small.

Previous theoretical work assessed the potential benefits of noise correlations violating the sign-rule [37, 41], and studied the interplay of noise and signal correlations in special cases with specific correlation structures [18, 19, 32, 36]. Previous decompositions of the mutual information [35, 52] suggested that variations of the noise correlations with the stimulus may be beneficial, with additionnal information encoded in these variations.

However, these results relied on a non-standard definition of noise correlations, making a direct comparison to our results intricate (see appendix D in [45]). Nonetheless, we considered the impact of such fluctuations within our framework, by relaxing the assumption of constant noise correlations in the second order derivation of the noise synergy (see Methods). The computation shows that these fluctuations can improve the noise synergy in two ways: by being large, and by being synchronized to the noise level *V*_n_(*θ*), also assumed to be stimlulus dependent. Our results thus extend and clarify previous theoretical work under a common information-theoretic framework.

Several studies have focused on the effect of noise correlations on the Fisher information [4, 18, 36, 39, 41]. While our main results are based on the mutual information, they equivalently apply to the Fisher information in the Gaussian case [33] (see Supplementary Appendix). To further test the robustness of our conclusions, we demonstrated that our results are model independent, and hold both for binary and Gaussian neurons. In addition, empirical results from the retinal recordings (Fig. 1J) were obtained without any approximation or model choice, and agree with the theory.

Also building on the Fisher information, another line of work [38, 53] suggested that noise correlations are detrimental when aligned to the signal direction in each point of response space. The structure of this type of noise correlations, called “differential” or “information-limiting” correlations, can be intuited from the definition of the Fisher information [38]. Although an in-depth discussion is beyond the scope of this paper, we have performed an additional numerical analysis (see Fig. S2) to demonstrate that information-limiting correlations become increasingly beneficial to the mutual information as their strength increases, while they are always detrimental to the Fisher information.

We validated our theoretical predictions experimentally on recordings of neurons from the retina. Applying our approach to data in sensory cortical areas where similar noise correlation structures have been observed [4, 5] could lead to new understanding of the role of noise correlations in sensory information processing. Another key open question is what stimulus ensembles most benefit from noise correlations, and where naturalistic stimuli stand in that regard. We have further shown that noise correlations benefit the encoding of high-frequency features of the stimulus, which correspond to fine-grained neural activity patterns. Extending these results to higher cortical areas would require understanding which features from the stimulus drive such activity patterns.

## IV. METHODS

### Covariance and correlation measures

The average responses of two neurons 1 and 2 are given as function of the stimulus *θ* by the tuning curves *µ*_1_(*θ*) = *r*_1 *θ*_ and *µ*_2_(*θ*) = *r*_2 *θ*_. Signal correlations are defined as *ρ*_s_ = Corr_*θ*_(*µ*_1_, *µ*_2_), and noise correlations as: *ρ*_n_(*θ*) = Corr(*r*_1_, *r*_2_ *θ*). The sum of these two coefficients does not have a simple interpretation in terms of total correlation or covariance, but we can also decompose the total correlation coefficient betweeen *r*_1_ and *r*_2_ as Corr(*r*_1_, *r*_2_) = *r*_s_ + *r*_n_, with 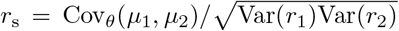, and 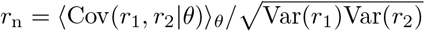.

### Pairwise analysis

#### Tuning curves

We consider a pair of neurons encoding an angle *θ*. The responses of the two neurons, *r*_1_ and *r*_2_, are assumed to be binary (spike or no spike in a 10 ms time window) and correlated. Their average responses *µ*_1_(*θ*) and *µ*_2_(*θ*) are given by Von Mises functions (Fig. 1A):

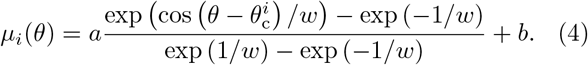

Signal correlations between the two neurons can be tuned by varying the distance between the center of the two tuning curves 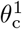 and 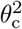. The tuning curve width *w* was set arbitrarily to 0.5, the amplitude *a* to 0.4 and the baseline *b* to 0.1. The strength of noise correlations is set to a constant of *θ, ρ*_n_(*θ*) = *ρ*_n_.

#### Small correlation expansion

When noise correlations *ρ*_*n*_ are constant, the noise synergy may be expanded as [45]:

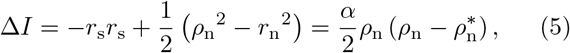

where the second equality highlights the dependency on *ρ*_*n*_. The critical 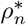 may be written as

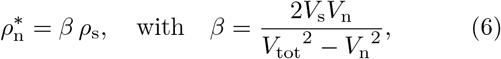

and the prefactor 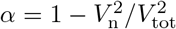, with the shorthands 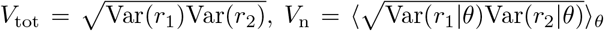 and 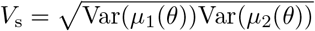 corresponding to measures of total, noise, and signal variances in the two cells.

By Cauchy-Schwartz we have:

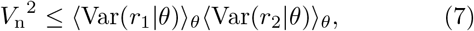

which entails

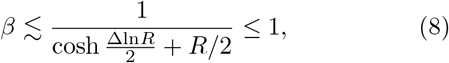

where 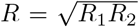 and Δln*R* = ln(*R*_1_*/R*_2_), with *R*_*i*_ = Var(*µ*_*i*_)*/* ⟨ Var(*r*_*i*_ *θ*) ⟩ _*θ*_ the signal-to-noise ratio of the cells. *R* measures the overall strength of signal-to-noise ratios, while *x* measures their dissimilarity. The last inequality implies that noise correlations are always beneficial for *ρ*_n_ *> ρ*_s_.

#### Varying noise correlations

When *ρ*_n_(*θ*) depends on *θ*, the noise synergy becomes [45]:

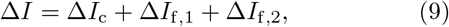

where Δ*I*_*c*_ is given by Eq. 5, and Δ*I*_f,1_ = (1*/*2)Var_*θ*_(*ρ*_n_(*θ*)) ≥ 0 accounts for the effect of fluctuations of *ρ*_n_(*θ*). Δ*I*_f,2_ is given by:

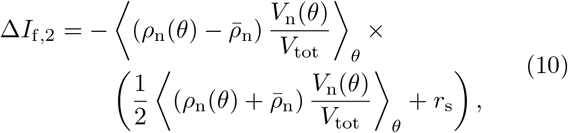

where 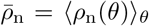. This contribution can be positive or negative, depending on how noise correlations *ρ*_n_(*θ*) co-vary with the noise variance of the pair *V*_n_(*θ*).

#### Gaussian case

To test the theory’s robustness to modeling choices, we also considered a continuous rather than binary neural response: *r*_*i*_ = *µ*_*i*_(*θ*) + *δr*_*i*_, where both *µ*_*i*_ and *δr*_*i*_ are Gaussian variables defined by their covariance matrices Σ_s,*ij*_ = Cov_*θ*_(*µ*_*i*_, *µ*_*j*_), and Σ_n,*ij*_ = ⟨ Cov(*r*_*i*_, *r*_*j*_ |*θ*) ⟩ _*θ*_. The noise synergy can be calculated through classic formulas for the entropy for Gaussian variables, yielding:

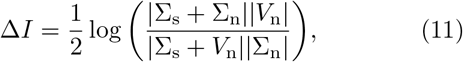

where |*X* | denotes the determinant of matrix *X*, and where *V*_n_ is the diagonal matrix containing the noise variances of the cells *V*_n,*ii*_ = Σ_n,*ii*_. Note that this formula is general for an arbitrary number of correlated neurons. In the pairwise case considered here matrices are of size 2*×*2. The condition for beneficial noise correlations Δ*I* ≥ 0 is satisfied for 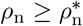,

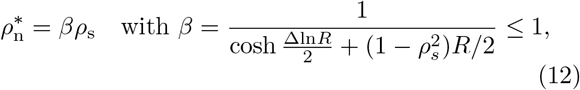

with which has a similar form as Eq. 8.

#### Noise synergy at constant noise entropy

Increasing noise correlations at constant *V*_n_ decreases the effective variability of the response, as measured by the noise entropy, *H*({*r*_1_, *r*_2_}|*θ*) = ln(2*πe*|Σ_n_|^1*/*2^), with 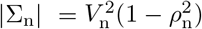 in the case two neurons with the same noise level. To correct for this effect we also computed Δ*I* at constant noise entropy, by rescaling the noise variances in the correlated and uncorrelated cases, *V*_n,c_ and *V*_n,u_, so that their resulting noise entropies are equal 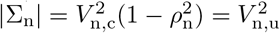.

The critical noise correlation at which Δ*I* ≥ 0 is then given by:

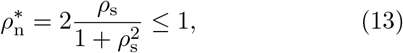

where the last inequality implies that strong enough noise correlations are always beneficial.

#### Retinal data

Retinal data were recorded ex-vivo from a rat retina using a microelectrode array [54] and sorted using SpyKING CIRCUS [46] to isolate single neuron spike trains. From the ensemble of single cells we could isolate a population of 32 OFF-*α* ganglion cells. Three stimuli movie with different spatio-temporal statistics were presented to the retina: a checkboard movie consisting of black and white checks changing color randomly at 40 Hz and repeated 79 times; a drifting grating movie consisting of black and white stripes of width 333 *µ*m moving in a fixed direction relatively to the retina, at speed 1 mm/s, and repeated 120 times; and finally a movie composed of 10 black disks jittering according to a Brownian motion on a white background, repeated 54 times.

### Gaussian population and spectral analysis

We consider a population of *n* neurons organized along a 1D lattice with constant interneuron spacing. Their mean response and noise are assumed to be Gaussian, with their noise and signal covariances given by an exponentially decaying function of their pairwise distances:

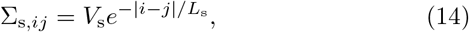

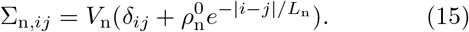

*V*_s_ and *V*_n_ are the signal and noise variance of the single cells. The parameter 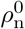 sets the strength of noise correlations such that nearest neighbors have noise correlation 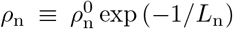. When *n* is large and boundary effects can be ignored, the system is invariant by translation and we can diagonalize Σ_s_ and Σ_n_ in the Fourrier basis 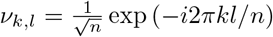. Denoting the spectra of Σ_s_ and Σ_n_ by *S*(*l/n*) and *N* (*l/n*), the expression of the noise synergy, Eq. 11, can then be written as a sum over modes:

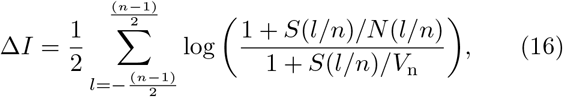

which simplifies in the *n* → ∞ limit to:

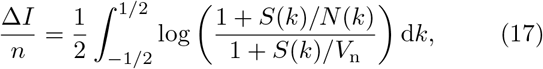

With

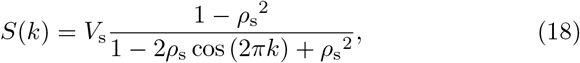

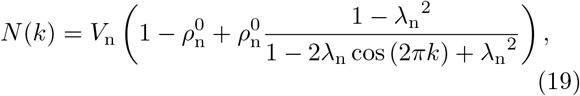

where *ρ*_s_ = exp (™1*/L*_s_) is the nearest-neighbors signal correlation, and *λ*_n_ = exp (− 1*/L*_n_). *k* is a wave vector interpretable as a spatial frequency in units of the system’s size, up to a 2*π* factor. Examining Eq. 17, we see that noise correlations are beneficial for frequencies for which *N* (*k*) ≤ *V*_n_, which happens for *k* ≥ *k*^*^ where *k*^*^ = (1*/*2*π*) arccos 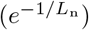.

In the low noise regime, *R* = *V*_s_*/V*_n_ ≫ 1, the noise synergy reduces to:

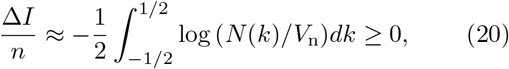

where the inequality stems from Jensen’s inequality, because − log is a convex function, and 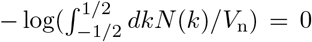. Therefore in that regime noise correlations are always beneficial.

In the high noise limit, *R* ≪ 1, the noise synergy becomes:

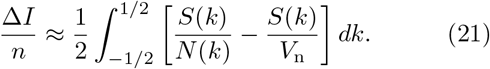

Computing this integral gives the critical noise correlation:

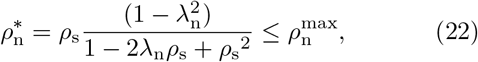

where 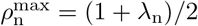 is the maximum possible value of *ρ*_n_ (ensuring that the noise spectrum *N* (*k*) is nonnegative for all *k*). The last inequality in Eq. 22 implies that there always exists a regime in which strong noise correlations are beneficial.

#### Non-identical neurons

To study the effect of nonhomogeneities among neurons, we considered the case where the signal variance of each cell is different, and drawn at random as 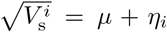, where *η*_*i*_ is normally distributed with zero mean and variance ν^2^. The noise synergy can be rewritten in the high noise regime (*R* ≪ 1) as:

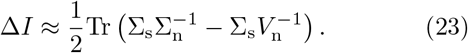

Averaging this expression over *η*_*i*_ yields:

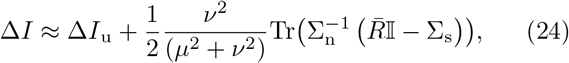

where Δ*I*_u_ is the noise synergy in a uniform population (with *V*_s_ = *µ*^2^ + ν^2^), and where the second term is always positive, with 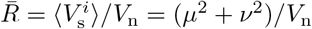.

Taking the continuous limit (*n* → ∞) in Eq. 24, similarly to the integral limit of Eq. 17, allows us to write the critical noise correlation 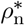 as:

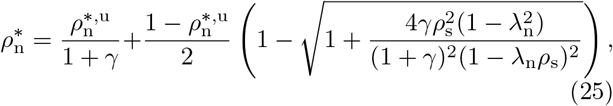

where *γ* = ν^2^*/µ*^2^ quantifies the relative magnitude of nonhomogeneities, and 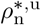 is the critical noise correlation value in a uniform population (Eq. 22). This modified critical noise correlation value is always smaller than in the uniform case, and scales linearly at leading order with the inhomogeneity parameter *γ*:

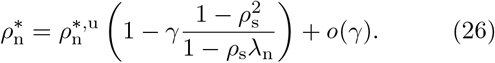

### Decoding analysis

#### Experimental and synthetic data

We presented a mouse retina with a stimulus consisting of a black and white random checkerboard flashed at 4Hz, each frame with a given spatial resolution (checks of sizes 12, 24, 36, 72 and 108 *µ*m). Retinal ganglion cell activity was recorded ex-vivo using a micro-electrode array and single neuron activity isolated via spike sorting using SpyKING CIRCUS [46]. We isolated a population of *N*_cells_ = 7 OFF-*α* retinal ganglion cells which presents strong noise correlations in their response [17]. The original recording contained a 15 *s* checkerboard movie repeated 90 times as well as 90 different 22.5 *s* long unrepeated movies.

We inferred a deep Generalized Linear Model (GLM) of the central cell among 7 from the experimental population (Fig. 4A), consisting of a stimulus-processing filter, and filters for the spiking history of the cell as well as its 6 neighbors (couplings). The stimulus-processing part of the model consisted of a deep neural network composed of two spatio-temporal convolutional layers followed by a readout layer. The whole model was fit to the data using the 2-step inference approach [47].

A synthetic population of 49 OFF-*α* ganglion cells was then constructed by arranging them on a triangular lattice of 7 by 7 points. Each cell responds according to the inferred GLM with translated receptive fields. Nearest neighbors were coupled with the average of the GLM couplings inferred between the central cell from the experiment and its neighbors.

To stimulate this synthetic population, we generated a synthetic stimulus ensemble from 550 regular black and white checker frames, each with a given check size ranging from 140 to 420 *µ*m (with increments of 28 *µ*m). Every checker of a given size was presented for 5 different regularly spaced offsets ranging from 0 to 224 *µ*m both in the horizontal and vertical directions, resulting in 25 different frames per size. To further ensure that the color of each pixel in the stimulus ensemble is black or white with equal probability, each checker frame also had its color-reversed version in the set, resulting in 50 different frames for a given check size. A single snippet from the synthetic stimulus ensemble consisted of a 250ms white frame followed by one of the 550 aforementioned checker frames.

We built a training, a vadliation, and testing set for the dependent and independent decoders by simulating the synthetic population for sets of 3750, 1250, and 5000 repetitions (respectively) of each synthetic stimulus snippets.

#### Decoders

The binary decoders are logistic regressors taking in the integrated response of the population over the *N*_*τ*_ = 5 past bins of 50 ms to predict the ongoing stimulus frame. The predicted stimulus at time *t* and repeat *k* is given by:

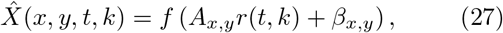

where *x, y* are the pixel indices along the two dimensions of the stimulus, *f* (*x*) = (1 +*e*^−*x*^)^−1^ is the sigmoidal function, *A*_*x*,*y*_ is a matrix of size (*N*_*τ*_, *N*_cells_), *r*(*t, k*) is a matrix of size (*N*_cells_, *N*_*τ*_) containing the spike history of the population at time *t* and repeat *k*, and *β*_*x*,*y*_ a pixelwise bias. Each decoder was trained by minimizing the average binary cross entropy (BCE) between predicted stimulus 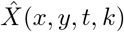 and the true stimulus *X*(*x, y, t*), (BCE(*x, y, t, k*))_*x*,*y*,*t*,*k*_, where

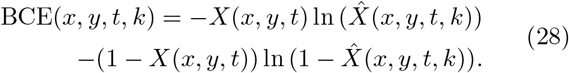

Training was done by stochastic gradient descent on the synthetic datasets using the training (3750 repetitions) and validation (1250 repetitions) sets. Optimization was done using stochastic gradient descent with momentum, with early stopping when the validation loss did not improve over 6 consecutive epochs. During that procedure the learning rate was divided by 4 whenever the validation loss did not improve for 3 consecutive epochs.

We probed the decoders’ abilities to decode features of different spatial scales by decoding the simulated responses of the synthetic population to the checker stimuli with varying check size from the testing set. Performances of the decoders were assessed by computing the mutual information between each pixel’s color *X* and it’s decoded value 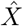, separately for the different sizes of checks. The noise synergy was then computed as the difference between the mutual information averaged over pixels for the dependent and independent decoders.

Error bars were computed as follows. We infered 10 deep GLMs on bootstraps of the original training set, obtained by re-sampling with replacement the simulusresponse pairs used for training. These 10 models were used to generate 10 surrogate training sets, from which 10 separate decoders were infered with noise correlations, and another 10 without noise correlations. Then synthetic test sets for the checker decoding task were generated from each of the 10 models, and the performance of each decoder computed separately with and without noise correlations, yielding 10 values of the mutual information, and 10 values of the noise synergy (both averaged over pixels). The error bars are the standard deviations of the resulting information, noise synergy and synergy-to-information ratios (i.e. relative noise synergy) over the 10 bootstraps.

## Data availibility

Part of the data utilized in this work have been published in previous studies. The remaining data and codes will be shared upon publication of this study. The code used to generate the synthetic data is available at https://github.com/gmahuas/noisecorr.

## Acknowledgements

We thank Stéphane Deny for help with the experimental data used in the paper and Matthew Chalk and Simone Azeglio for useful discussions. This work was supported by ANR grants n. ANR-22-CE37-0023 “LOCONNECT” and n. ANR-21-CE37-0024 NatNetNoise, and by IHU FOReSIGHT (ANR-18-IAHU-01) and by Sorbonne Center for Artificial Intelligence-Sorbonne University-IDEX SUPER 11-IDEX-0004. This work was also supported by the Bettencourt Schueller Foundation.

---

**FIG. S1:**
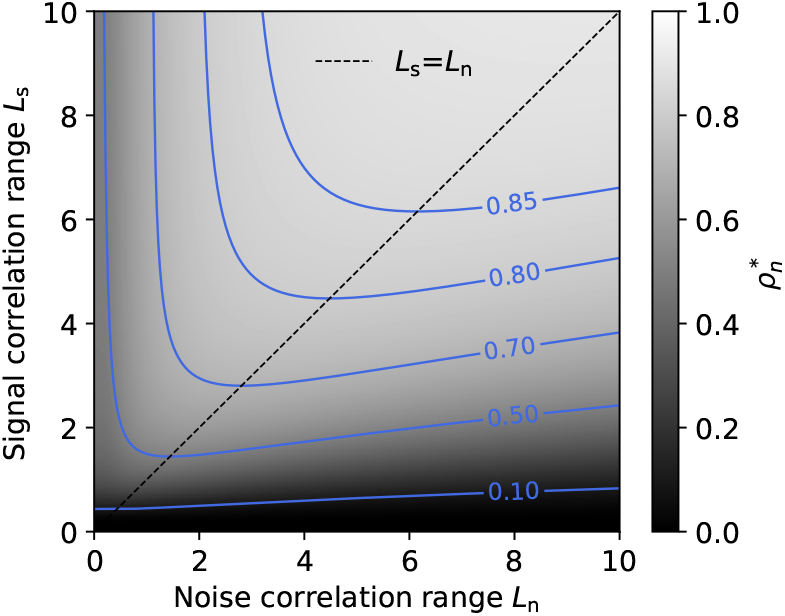
**Behavior of** 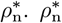 changes non-monotically with the signal *L*_s_ and noise *L*_n_ correlation ranges, and is concave with respect to these parameters. The maximum value of 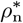 at a given *L*_s_ is achieved when *L*_n_ = *L*_s_.

## Supplementary information

### Strong noise correlation and Fisher information

We consider a pair of neurons encoding an arbitrary variable *θ*. The responses *r* of these neurons is assumed to be Gaussian of mean *µ*(*θ*) and covariance matrix Σ_n_), where *µ* (*θ*) are the tuning curves. In this context the Fisher information is defined as:

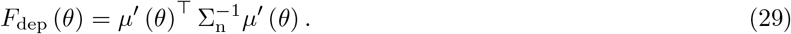

Expanding this expression for a pair of neurons with equal noise variance, 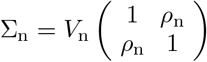, yields:

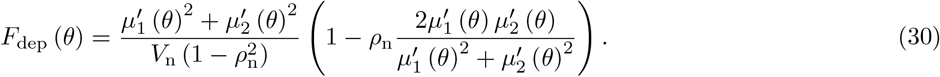

In the absence of noise correlations (*ρ*_n_ = 0), the Fisher information simplifies to:

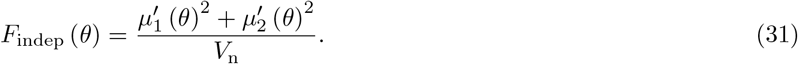

To quantify the overall Fisher improvement we introduce the quantity Δ*R* = (*F*_dep_(*θ*)*/F*_indep_(*θ*) ™ 1) _*θ*_. Defining 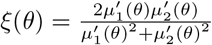, the Fisher improvement becomes:

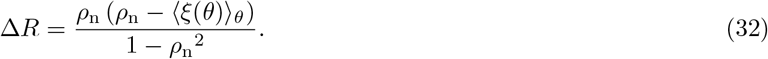

Therefore, strong positive noise correlations will benefit the Fisher information whenever they exceed the critical value 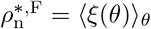.

**FIG. S2:**
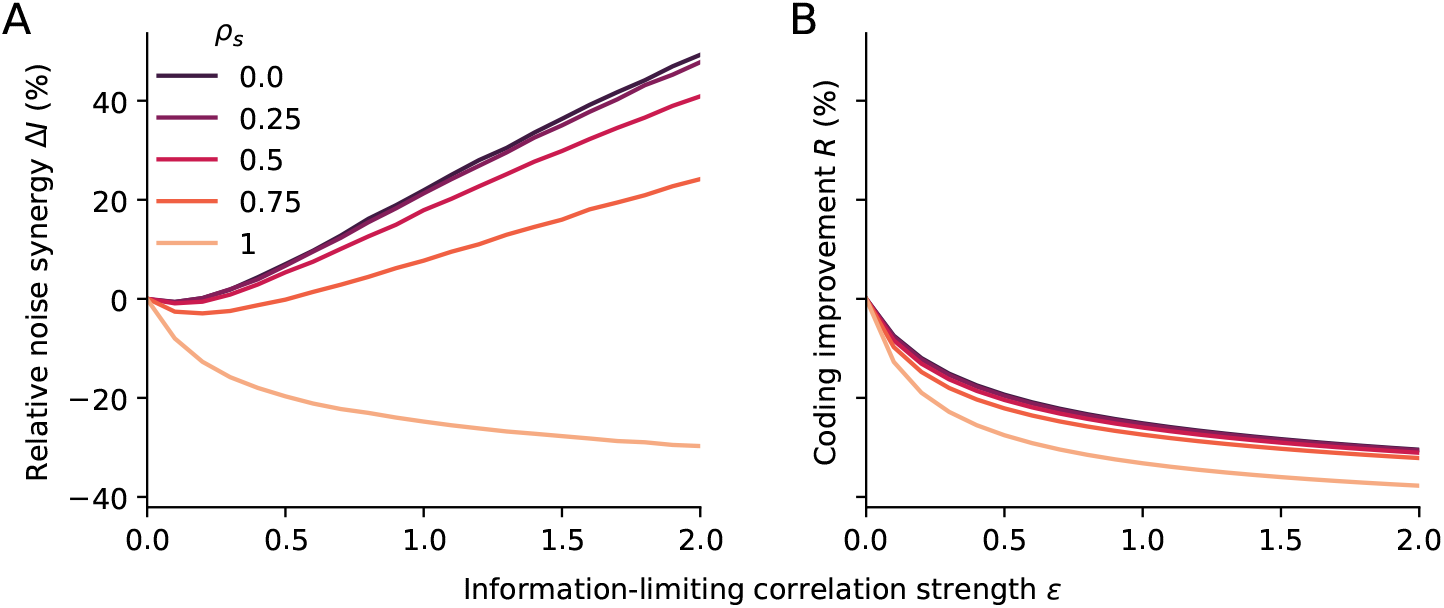
Impact of information limiting correlations on stimulus information. An angular stimulus *θ* is encoded by a pair of neurons characterized by Von Mises tuning curves (with parameters *a* = 40, *b* = 10, and *w* = 5). Their response is Gaussian of means *µ*_1_(*θ*) and *µ*_2_(*θ*). Information-limiting correlations are defined by a covariance of the form: Σ_n_(*θ*) = *V*_0_II + *Eµ*^*t*^(*θ*)*µ*^*t*^(*θ*)^*T*^, were *E* controls their strength, and where we set *V*_0_ = *V*_s_*/*2. Note that Σ_n_ now depends on *θ*. **A**. Relative noise synergy Δ*I* = *I*_dep_*/I*_indep_ − 1 as a function of *E*, where *I*_dep_ and *I*_indep_ quantify the mutual information with and without (off diagonal terms of Σ_n_ set to 0) noise correlations, for different levels of signal correlation *ρ*_s_. Mutual information was computed via Monte-Carlo integration. Information-limiting noise correlations become beneficial to the mutual information if they are strong enough, except when cells are perfectly signal-correlated. **B**. By contrast, information-limiting correlations are always detrimental to the Fisher information 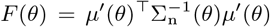. The relative Fisher improvement Δ*R* = ((*F*_dep_(*θ*)*/F*_indep_(*θ*)−1))_*θ*_, where *F*_dep_(*θ*) and *F*_indep_(*θ*) denote the Fisher information with and without noise correlations, is always negative and decreases with *ϵ* and *ρ*_*s*_.

